# *Shank3* establishes AMPA receptor subunit composition at cerebellar mossy fiber-granule cell synapses and shapes regional microglia activation

**DOI:** 10.1101/2025.08.01.668222

**Authors:** Rajaram Kshetri, Ben D Richardson

## Abstract

Mutations in *Shank3* are the primary genetic cause of Phelan-McDermid Syndrome (PMS), a neurodevelopmental disorder frequently comorbid with autism spectrum disorder (ASD). As a key scaffolding protein in the postsynaptic site, SHANK3 is critical for excitatory glutamatergic synapse function by interacting with AMPARs, NMDARs, and mGluRs. While *Shank3* deficiency has been extensively studied in forebrain regions, its role in the cerebellum, a brain area increasingly implicated in ASD pathobiology, remains comparatively underexplored. Cerebellar granule cells (CGCs) exhibit high *Shank3* expression. However, its role in cerebellar glutamatergic synapses is poorly understood. This study aims to investigate how *Shank3* loss affects mossy fiber-CGC glutamatergic synaptic function.

Whole-cell patch clamp electrophysiological recordings from CGCs in *ex vivo* cerebellar brain slices from adult (4-6 months old) wild type (WT) and homozygous *Shank3*^Δ*ex4-22*^ KO were performed to record miniature, evoked, and glutamate uncaged responses. Similarly, the current-voltage (I-V) relationship was analyzed with intracellular spermine and pharmacological validation of calcium-permeable AMPARs (CP-AMPARs) was done by IEM-1460. Immunofluorescence staining was performed for microglia using IBA1 labeling.

We found a significant increase in mEPSC amplitude and AMPAR-mediated response to glutamate uncaging, which indicates that the loss of *Shank3* enhances postsynaptic AMPAR function. Furthermore, the KO group showed faster AMPAR decay kinetics, inward rectification, and increased sensitivity to IEM-1460, suggesting that a high proportion of CP-AMPARs with distinct biophysical properties are present at the MF-CGC synapse. Furthermore, KO mice showed less ramified microglia suggesting the possible presence of activated microglia in the cerebellar cortex.

Together, these findings highlight a critical role of *Shank3* in maintaining the balance between CP- and CI-AMPARs at the MF-CGC synapse, which is essential for synapse maturation and proper cerebellar circuitry function. Dysregulation of this balance, with possible presence of activated microglia in the cerebellum, may underscore cerebellar-related behavioral deficits in *Shank3* KO mice and may suggest a potential mechanism contributing to ASD pathophysiology.

## Background

Autism spectrum disorder (ASD) is a complex neurodevelopmental disorder with a heterogeneous group of symptoms typically diagnosed in early childhood (Jung & Park, 2022; H. Kim et al., 2016; K. Wang et al., 2017). ASD is characterized by a group of core behavioral symptoms which include impaired social communication, restricted interactions, interests or activities, and repetitive behaviors. (Berkel et al., 2018; *Diagnostic and Statistical Manual of Mental Disorders: DSM-5^TM^*, 2013; Gandhi & Lee, 2021; H. Kim et al., 2016; Lord et al., 2020; K. Wang et al., 2017). Disrupted synaptic structure and function has been often proposed as a converging mechanism underlying the diverse phenotypic manifestations of ASD. The identification of numerous ASD-associated genes that encode proteins integral to synaptic biology has led to the concepts of “synaptic hypothesis” or “synaptopathy” (Bauman & Kemper, 2005; Sahin & Sur, 2015; Won et al., 2013).

*SHANK3* is widely recognized as a prominent candidate gene strongly associated with ASD (Betancur & Buxbaum, 2013; Hulbert & Jiang, 2017). Along with being the cause of Phelan-McDermid Syndrome (Betancur & Buxbaum, 2013), a mutation or deletion affecting *SHANK3* accounts for approximately 1-2% of all ASD cases, making it one of the prominent monogenic causes of ASD with a high penetrance for ASD phenotypes (Jiang & Ehlers, 2013; Soorya et al., 2013). Although classically considered as a regulator of postsynaptic glutamate receptors, Shank3 plays an important role in facilitating protein-protein interactions between neurotransmitter receptors, ion channels, intracellular cytoskeleton components, and signal transduction pathways (Schmeisser & Verpelli, 2016; Wan et al., 2021). Shank3 protein is expressed as different isoforms in the cortex, hippocampus, striatum, thalamus, and cerebellum (Monteiro & Feng, 2017; Peça et al., 2011) with *Shank3a, b,* and *e* highly expressed in the striatum and *Shank3c* and *d* in the cerebellum (X. Wang et al., 2014). Deletion of *Shank3* affects synaptic function in striatum (Peça et al., 2011; Yoo et al., 2018), hippocampus (Bey et al., 2018; Bozdagi et al., 2010; Kouser et al., 2013), thalamus (B. Guo et al., 2024), and cortex (B. Guo et al., 2019; Yoo et al., 2019).

Considerable progress has been made in explaining the neurobiological basis of synaptic dysfunction, with most studies reporting alterations in excitatory glutamatergic synaptic transmission in *Shank3* mutant mice (Berg et al., 2018; Bozdagi et al., 2010; Jiang & Ehlers, 2013; Sala et al., 2015; Wan et al., 2021). However, heterogeneity is observed depending on the specific genetic manipulation, brain region, and developmental stage examined (Jung & Park, 2022; Zhang et al., 2023). *Shank3*^Δ*e4–22*^ mouse model has shown alterations in physiological processes, particularly synaptic transmission and plasticity in the striatum (Bey et al., 2018; Drapeau et al., 2018; X. Wang et al., 2016). *Shank3* deletion is reported to increase basic cellular excitability (Bozdagi et al., 2010; X. Wang et al., 2011) and decrease the frequency of spontaneous excitatory postsynaptic currents (sEPSCs), as well as impair long-term depression (LTD) of medium spiny neurons (MSNs) in the striatum (X. Wang et al., 2011). Similarly, conditional KO of *Shank3* exons 4-22 in neocortical excitatory neurons has shown an increase in the NMDA/AMPA ratio in CA1 neurons of the hippocampus (Bozdagi et al., 2010).

Over the past two decades, many studies have highlighted the cerebellum as a key brain region implicated in ASD pathogenesis (Becker & Stoodley, 2013; Fatemi et al., 2012; Hampson & Blatt, 2015; Mosconi et al., 2015; Tsai, 2016; S. S. H. Wang et al., 2014). Several experiments using animal models with ASD-related genes (*Tsc1, Shank2, Mecp2*) have been conducted to investigate the role of the cerebellum, primarily focusing on Purkinje cells (PCs) (Achilly et al., 2021; S. Ha et al., 2016; Kloth et al., 2015, 2015; Stoodley et al., 2017; Tsai et al., 2012). These studies were critical in establishing a role for ASD-linked genes specifically in PCs to shape behaviors outside of the motor domain. However, gene/mRNA expression data (Furuichi et al., 2011; Lein et al., 2006; Lonsdale et al., 2013; Menashe et al., 2013; Sato et al., 2008; Satterstrom et al., 2020) suggest a role for ASD-linked genes in the CGCs that integrate all multisensory input entering the cerebellar cortex to then determine the firing behavior of PCs. Alteration of some ASD-linked genes (*Chd8, Ib2*) expressed by CGCs has shown motor dysfunction as well as alterations in CGC excitability and synaptic functions (Kawamura et al., 2021; Soda et al., 2019). Overall, the delineation of mechanisms by which ASD-linked genes affect the physiology of cerebellar cortical neurons, especially CGCs, and how such genes in the cerebellum contribute to motor and non-motor processes is lacking.

Additionally, some studies have shown the potential involvement of glial cells (astrocytes and microglia) in contributing to the neuroinflammatory profile observed in ASD (Bailey et al., 1998; Vargas et al., 2005; Ahmad et al., 2017; Matta et al., 2019). Postmortem studies from individuals diagnosed with ASD have frequently revealed neuroinflammation, especially within the cerebellum (Vargas et al., 2005). Furthermore, neuropathological investigation of postmortem brain tissue from individuals with ASD has consistently demonstrated alterations in microglial characteristics (Andoh et al., 2019; Fan et al., 2023; Hu et al., 2022; Hughes et al., 2023; Matta et al., 2019; Petrelli et al., 2016; Xiong et al., 2023, 2023). These studies reported an increased microglial number and a morphology suggestive of an activated state, in multiple brain regions, including the frontal lobes and cerebellum. Despite this compelling evidence of altered microglial states and increased inflammatory markers in ASD, a fundamental question persists regarding whether this observed neuroinflammation and glial activation represents a primary factor in ASD pathogenesis or is a secondary consequence of other underlying pathological processes.

In our previous study, we found that the *Shank3*^Δ*e4–22*^ mouse demonstrated behavioral impairments, mainly affecting motor function, anxiety, and repetitive behaviors, especially in adult mice (Kshetri et al., 2024), which was paralleled by AMPAR-mediated response augmentation. Despite the high expression of Shank3 in CGCs and its impact on AMPAR-mediated EPSC size in late development, how Shank3 affects AMPAR function and the consequences of these specific changes are unclear. In this study, we aimed to investigate how the loss of *Shank3* alters glutamtatergic synaptic function at the MF-CGC synapse and how it may affect local supporting cell morphology and CGC density in the cerebellar cortex.

## Materials & Methods

### Animals

All animal procedures were performed in accordance with the protocols approved by the Institutional Animal Care and Use Committee (IACUC) at the Southern Illinois University School of Medicine (Protocol #2023-129). All animals were group-housed, provided a normal chow diet, and maintained on the reverse a 12-hour light-dark cycle to facilitate behavioral experiments during their active (dark) phase. Offspring genotypes were determined by Transnetyx (Cordova, TN) using ear punch or tail biopsies. *Shank3*^Δ*ex4–22*^ mice lack exons 4-22 of the *Shank3* gene and hence lack expression of all major SHANK3 protein isoforms A to F (Bozdagi et al., 2010; Drapeau et al., 2018; X. Wang et al., 2014). A breeder pair of heterozygous *Shank3*^Δ*ex4–22*^ mice maintained on C57BL/6NJ genetic background (Drapeau et al., 2018) (JAX strain #: 032169) were acquired from the Jackson Laboratory (Bar Harbor, ME) and bred in-house to generate wildtype (+/+, WT) and homozygous knockout (-/-, KO) mice. All electrophysiology and histological experiments used only adult (4-6 months) WT and KO mice.

### Cerebellar slice electrophysiology

To prepare acute brain slices for recording from CGCs, adult (4-6 months) mice were anesthetized with 3% isoflurane, and cardiac perfusion was done with artificial cerebrospinal fluid (ACSF) containing 1 mM kynurenic acid. Then, the brain was rapidly removed and placed in an ice-cold sucrose slicing solution. The modified form of sucrose slicing solution and recovery solution mentioned in Chabrol et al., 2015 was used in this study. This solution contained the following components (in mM): 2.5 KCl, 0.5 CaCl_2_, 4 MgCl_2_, 1.25 NaH_2_PO_4_, 24 NaHCO_3_, 25 glucose, 230 sucrose, and 1 kynurenic acid. The brain was then mounted on a holder and encased in agar and sliced into a parasagittal section (250 μm) using a Compresstome VF-200 (Precisionary Instruments). The cerebellar slices were then transferred to a recovery solution that included the following components (in mM): 85 NaCl, 2.5 KCl, 0.5 CaCl_2_, 4 MgCl_2_, 1.25 NaH_2_PO_4_, 24 NaHCO_3_, 25 glucose, 75 sucrose, and 1 kynurenic acid maintained at 32 °C (Chabrol et al., 2015). After 30 minutes of recovery, cerebellar slices were transferred to room temperature ACSF containing (in mM): 124 NaCl, 26 NaHCO_3_, 1 NaH_2_PO_4_, 2.5 KCl, 2 MgCl_2_, 10 D-glucose, and 2.5 CaCl_2_. All solutions were saturated with 95% O_2_ and 5% CO_2_, had a pH of 7.3–7.4, and osmolarity of 300–310 mOsm. Slices were transferred to a custom recording chamber on an upright Olympus BX51WI microscope, and CGCs in the internal granule cell layer in lobules 4–5 were visualized with a 60X water-immersion objective using infrared differential interference contrast. ACSF was continuously perfused into the chamber at a rate of 3–5 ml/min, maintained at 32–34 °C.

Whole-cell voltage-clamp recordings of visually identified CGCs were made using borosilicate patch pipettes (1.5 mm OD/0.86 mm ID) pulled with a P-1000 micropipette puller (Sutter Instruments) to have a tip resistance of (5–8 MΩ) when filled with CsCl-based internal solution (E_Cl_ = 0 mV) that contained (in mM): 130 CsCl, 4 NaCl, 0.5 CaCl_2_, 10 HEPES, 5 EGTA, 4 Mg-ATP, 0.5 Na-GTP, and 5 QX314 with pH adjusted to 7.2–7.3 with CsOH and an osmolarity of 280–290 mOsm (Kaplan et al., 2016; Richardson & Rossi, 2017). Whole-cell patch-clamp recordings were acquired with a Multiclamp 700B amplifier (Molecular Devices) and sampled at 20 kHz (10 kHz low pass filter) with a Digidata 1440 A (Molecular Devices). Following the formation of a gigaseal (> 1GΩ), the whole-cell configuration was produced by the application of rapid negative pressure to the pipette. Whole-cell membrane properties were determined by applying a 10 mV hyperpolarizing voltage step from the initial holding potential (-60 mV) in voltage-clamp mode. Whole cell recordings from CGCs had a series resistance of 20 ± 5 MΩ and recordings with variation in series resistance of greater than 20% throughout the recording were discarded.

### Miniature excitatory postsynaptic currents (mEPSCs)

To isolate miniature excitatory postsynaptic currents (mEPSCs), CGCs were voltage-clamped at -60 mV and the GABA_A_ receptor antagonist, gabazine (SR95531; 10 μM; Tocris Bioscience, catalog# 1262) was present in the ACSF. To isolate mEPSCs, CGCs were voltage-clamped at -70 mV with gabazine and the voltage-gated sodium channel blocker tetrodotoxin (TTX; 0.5 µM; Tocris Bioscience) was included in the bathing ACSF.

#### Data analysis

The acquired raw trace was filtered offline with a 2 kHz lowpass gaussian filter in Clampfit 11.2 (Molecular Devices). Then, inward transient mEPSCs with a fast rise and exponential decay were analyzed over a 3–5 min period with Easy Electrophysiology Software (v2.6.0) by first-pass automatic threshold detection followed by manual inspection of events. All events from each CGC were used to construct a cumulative distribution histogram for amplitude (1 pA bin size) or inter-event interval (IEI, 100 ms bin size). Event amplitude and IEI were averaged for each cell to generate group averages and for statistical comparisons between genotypes.

### Evoked excitatory postsynaptic currents (eEPSCs)

A bipolar tungsten electrode (0.5 MΩ; World Precision Instruments) was placed in the central fiber bundle within the cerebellar lamina of lobe 4 & 5 to stimulate the presynaptic mossy-fiber terminals. Stimulation was performed using a range of stimulus intensities from 50 to 300 µA in 50 µA increments. The stimulation frequency was set at 0.2 Hz, with paired-pulse stimulation using interstimulus intervals of 20 ms. To calculate the input-output curve, electrical stimulation intensities ranging from 50-300 µA were used, while CGCs were held at a membrane potential of -60 mV. A stimulation intensity of 100 µA was used to calculate the peak AMPA, peak NMDA, paired-pulse ratio (PPR), and AMPA/NMDA ratio. During presynaptic terminal stimulation, the CGCs were initially maintained at -60 mV in the presence of gabazine (10 µM) to record the AMPAR-mediated response. Subsequently, the holding potential of CGCs changed to +40 mV to record the combined AMPA and NMDAR-mediated response.

#### Data analysis

At least 10-15 non-failure events were selected and digitally averaged for each stimulus recording. The averaged trace was then filtered using a 2 kHz Gaussian filter in Clampfit 11.2 software (Molecular Devices) to obtain an average response for AMPA- and NMDAR-mediated currents. The amplitudes of AMPAR and NMDAR currents were calculated by placing a 1 ms baseline window before the stimulation. The AMPA/NMDA ratio was determined by calculating the ratio of the peak AMPA current at -60 mV and NMDA current value after 15 ms of stimulation at +40 mV (Giza et al., 2010).

To calculate AMPAR-mediated decay, the peak-to-baseline decay phase of the resulting current trace was fitted by the double exponential function: I = A_1_e ^−t/τ1^ + A_2_e ^−t/τ2^ and the weighted decay constant was calculated using the following formula where, τ_W_ = weighted decay constant, τ_1_ = slow decay time constant, τ_2_= fast decay time constant, A_1_ = slow amplitude component, and A_2_ = fast amplitude component (Richardson et al., 2013; Schofield & Huguenard, 2007):

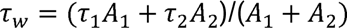

### Rubi-Glutamate uncaging

For the glutamate photo-uncaging experiment, we used a caged glutamate compound called Ruthenium-bipyridine-trimethylphosphine-Glutamate (RuBi-Glutamate). This compound can be excited by visible wavelengths and releases glutamate upon one- or two-photon excitation (Fino et al., 2009). The CGC was maintained at a holding potential of -70 mV, and photo-evoked EPSCs were recorded in the whole-cell configuration. Focal stimulation of CGCs was achieved by applying 470 nm blue LED light within a 50 × 50 µm² gridded area over the cerebellar cortex. To facilitate the photo-release of glutamate, the Polygon 1000 pattern Illuminator (Mightex) was used. This experiment was conducted in the presence of 50 µM RuBi-Glutamate, chosen as the optimal concentration based on reliable response using the stimulation protocol. Glutamate photo-release was achieved using a light stimulation duration of 50 ms at a frequency of 0.05 Hz. To record the combined AMPA + NMDA current evoked by photo-stimulation, the CGC was held at -70 mV, and the cerebellum slice was perfused with ACSF containing RuBi-glutamate, 10 µM gabazine, and 0.5 µM tetrodotoxin (TTX; Tocris Bioscience, catalog # 1069) in absence of Mg^2+^. NMDA current was pharmacologically dissected by subsequent supplementation with 10 µM NBQX in the bath. The NMDA current was further validated by subsequent bath application of 50 µM AP5 and 50 µM 7-Chlorokynurenic acid. After the drugs were introduced into the bath, at least 2-3 min were allowed for proper diffusion, and then the current responses were recorded.

#### Data analysis

At least five traces with a plateau response were selected, digitally averaged, and filtered using a 200 Hz Gaussian filter in Clampfit 11.2 software (Molecular Devices) to obtain an average response for each current type, i.e., AMPA + NMDA, AMPA, and NMDA. The peak amplitude of these currents was calculated by placing a 1 ms baseline window before the light was turned on for uncaging. The AMPA trace was obtained by digitally subtracting the NMDA trace from the AMPA + NMDA trace in Clampfit 11.2. The AMPA/NMDA ratio was calculated by dividing the AMPA-mediated peak amplitude by the NMDA-mediated peak amplitude. The current density was calculated by dividing the peak amplitude currents by the cell capacitance.

### Inward rectification of eEPSCs

In the same eEPSC recording mode, a patch pipette filled with a CsCl-based internal solution supplemented with 100 µM spermine (Tocris Bioscience, catalog# 958) was used to perform the inward rectification experiment. First, CGC was held at -60 mV, and single-pulse stimulation of 100 µA at a frequency of 0.2 Hz was applied in the presence of the pharmacological blockers 10 µM gabazine to record AMAPR-mediated eEPSC. Then, the holding potential was changed from -60 to +60 mV with successive increments of 20 mV, and the bath was supplemented with additional NMDAR blockers: D-AP5 (50 µM; Tocris Bioscience, catalog # 0106) and 7-Chlorokynurenic acid (50 µM; Tocris Bioscience, catalog# 3697) to record the AMPAR peak current at positive potentials.

#### Data analysis

At least 10-15 traces were selected and digitally averaged for each stimulus recording, then the averaged trace was then filtered using a 2 kHz Gaussian filter in Clampfit 11.2 software (Molecular Devices) to obtain an average response for AMPAR-mediated currents. The amplitude of AMPAR currents was calculated by placing a 1 ms baseline window before the stimulation. A normalized current-voltage (I-V) curve was generated by normalizing the average peak AMPAR current values at each holding potential to the maximum response at V_h_ = -60 mV. The rectification index was calculated by taking the ratio of the peak AMPAR current at +60 mV to the AMPAR current at -60 mV. **Pharmacological assessment of IEM-1460 sensitivity**

In the same eEPSC experimental setup using CsCl-based internal solution, the AMPAR-mediated response was recorded at -60 mV after stimulating with single-pulse stimulation of 100 µA at a frequency of 0.2 Hz in the presence of gabazine (10 μM; Tocris Bioscience, catalog# 1262) followed by the supplemental application of selective calcium-permeable AMPAR blocker IEM-1460 (100 µM; Tocris Bioscience, catalog# 1636). The baseline was recorded for the first 5 min in the presence of gabazine, which was followed by the additional supplementation of IEM-1460 and recording for an additional 15 min.

#### Data analysis

The average value of 12 traces that were recorded every minute were selected to obtain the average peak AMPAR-mediated response. The amplitude of AMPAR currents was calculated by placing a 1 ms baseline window before the stimulation. A normalized current-voltage (I-V) curve was generated by normalizing the average peak AMPAR current values at each holding potential to the maximum response at V_h_ = -60 mV. The average responses each minute after the application of IEM-1460 were normalized to their respective average value of their baseline recorded for first 5 min. The percentage of baseline response was calculated by calculating the average percentage of response in last 3 min in IEM-1460 compared to the average baseline response.

### Immunofluorescence

For all immunofluorescence studies, mice were anesthetized using isoflurane (3-5%) and then transcardially perfused with 1X phosphate-buffered saline (PBS), followed by 4% formaldehyde diluted in 1X PBS. Then, brains were removed and post-fixed for 48–72 h in 4% formaldehyde, and placed in sucrose solution gradients until they sank to the bottom. Following the sucrose gradient, brain samples were snap-frozen in dry ice before embedded in optimal cutting temperature (OCT) (Tissue-Tek; Sakura Finetek, CA, catalog# 4583). Sagittal whole-brain slices (30 μm thick) containing cerebellar vermis were prepared on a cryostat (Leica CM 1850) at -25 °C. Subsequently, the brain slices were washed in 1X PBS and then stored at -20 °C in a cryoprotecting solution until the start of immunofluorescence studies. All immunofluorescence staining procedures were performed using the free-floating method.

### IBA1 staining

The brain slices were placed in sodium borohydride (1 mg/ml in PBS) for 30 min, replacing the fresh solution every 10 min at room temperature. The brain slices were then permeabilized in 0.25% Triton X-100 in PBS for 30 min, followed by blocking in IHC/ICC Blocking Buffer (eBiosciences) containing 0.25% Triton X-100 for 1 h. Subsequently, the slices were incubated in primary antibody rabbit IgG anti-IBA1 (1:500; Wako, catalog# 019-19741) overnight at 4 °C. Slices were then washed with PBS and incubated in secondary antibody goat IgG anti-rabbit Alexa Fluor 488 (1:1000; Thermo Fisher Scientific, catalog# A11008). The nuclei were counterstained with Hoechst 33342 (1:2000, Thermo Fisher Scientific, catalog# H3570). Finally, tissue sections were washed and transferred to glass slides and mounted with Prolong Gold (Invitrogen). Imaging of microglia in the inner granule cell layer of cerebellar lobe 4 &5 from the stained brain slices was performed by capturing Z-stack images containing 7 optical slices at an interval of 1 µm, using a 40X oil immersion objective (numerical aperture 1.3) on a confocal microscopy system (Zeiss LSM800) at a resolution of 1,024 × 1,024 pixels. *Data analysis:* The orthogonal projections of Z-stack images were generated using ZEN Lite software (v 3.7 edition; Zeiss). The total surface area covered by individual microglia was determined using Imaris software (v10.2). The total number of microglia was assessed by counting the number of IBA1-positive microglial cell bodies in each orthogonal projection image. Then, the proportion of microglia of different surface area measurements was quantified. Total 5-6 images/genotype were analyzed from 3-5 animals/genotype.

### Statistical analysis

For analyzing electrophysiology and immunofluorescence data, the normality of data was tested using Shapiro-Wilk test. An unpaired t-test was used to compare all the parameters between WT and KO mice within age groups using GraphPad Prism 10.4.2. In all experiments, data values are reported as mean ± standard error (SEM) with individual markers representing the value for each observation, which is the cell (n) for electrophysiology assays, and the image (n) for confocal analysis. All statistical comparisons with p < 0.05 were considered significant. Statistical results are provided in **Supplementary Table 1**.

## Results

### Loss of *Shank3* alters postsynaptic AMPAR function at MF-CGC synapse

To determine the cause of augmented MF-CGC synaptic function we identified previously (Kshetri et al., 2024), we investigated stochastic spontaneous neurotransmitter release and changes in synaptic function at the MF-CGC synapse due to the loss of *Shank3.* First, quantal glutamatergic miniature excitatory postsynaptic currents (mEPSCs) resulting from single vesicle release from presynaptic mossy fiber terminals were recorded from CGCs at a holding potential of -70 mV in the presence of 10 µM gabazine and 0.5 µM TTX (**Figure 1A**). Since TTX blocks voltage-gated sodium channels, which are essential for generating action potentials and coordinated multivesicular release, mEPSC recordings provide an assessment of synaptic responses to individual vesicles/single quanta by eliminating the possibility of multivesicular release.

**Figure 1:**
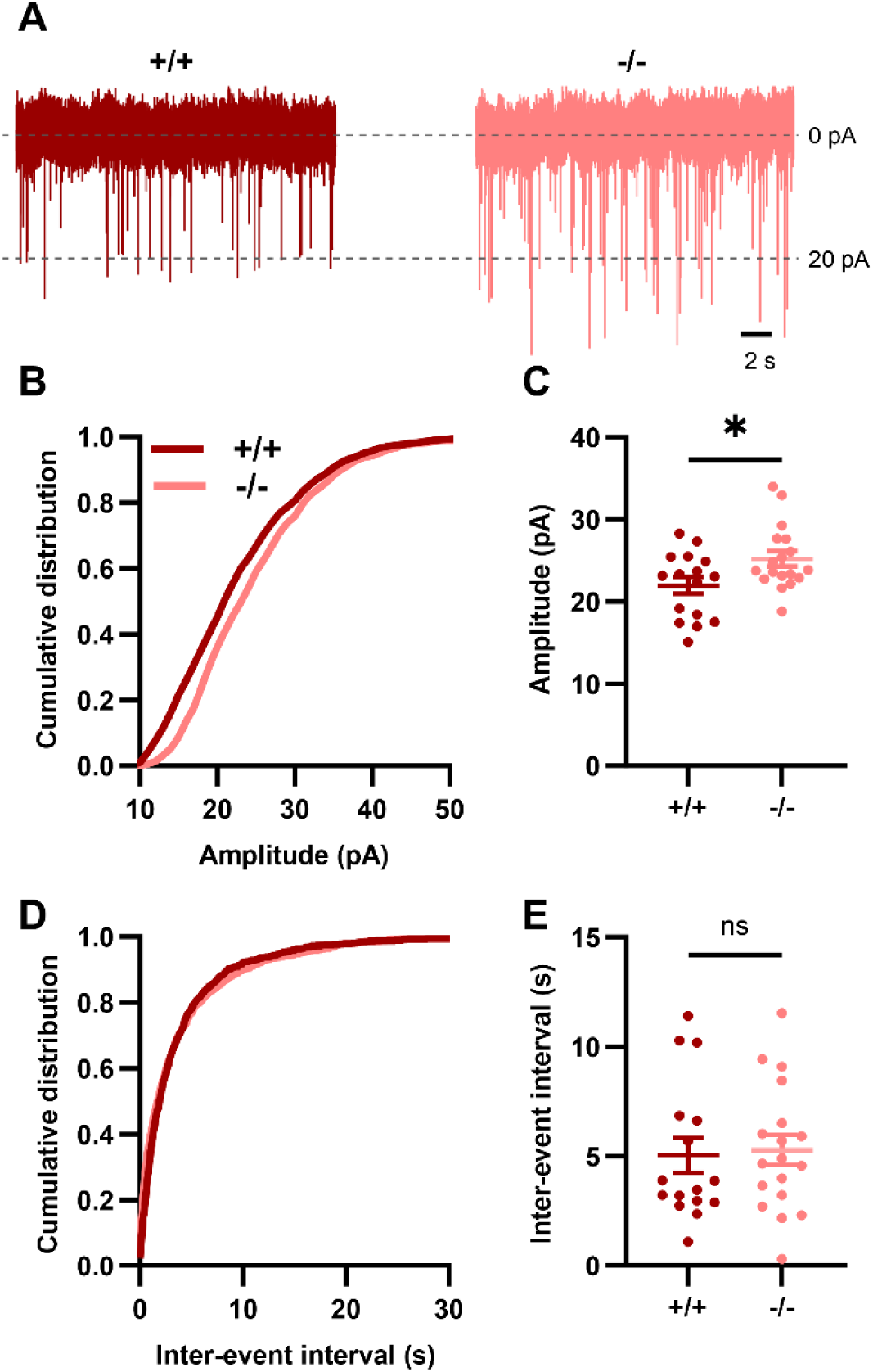
Increased mEPSC amplitude in CGCs of adult *Shank3* KO mice. (**A**) Representative traces of mEPSC recorded from CGC in WT (+/+, red) and *Shank3* KO (-/-, light red) mice in the presence of 10 μM gabazine and 0.5 μM TTX. (**B, D**) Cumulative distribution histograms of mEPSC amplitudes (**B**) and interevent intervals (**D**) for all events from WT and KO groups. (**C, E**) Individual average data points from each cell (circles) and group mean ± SEM (bars) for mEPSC amplitudes (**C**) and IEIs (**E**). WT: n = 16 cells from N = 10 mice, 1419 events; *Shank3* KO: n = 18 cells from N = 10 mice, 1391 events. Statistical significance was determined using an unpaired t-test for mEPSC amplitude (**B**) and a Mann-Whitney U test for IEI (**C**). *p < 0.05 indicates a significant difference between genotypes; ns: not significant.

We also assessed tonic inhibition current in CGCs by applying 10 µM gabazine, but we did not find a significant difference between WT and KO mice (t(19) = 1.179, p = 0.25). Similar to our previous observation in sEPSCs (Kshetri et al., 2024), cumulative distribution of mEPSC amplitudes was shifted toward higher values, indicating a change in postsynaptic AMPAR function in CGCs (**Figure 1B**). The average peak amplitude of mEPSCs was significantly larger in the *Shank3* KO group compared to the WT group (**Figure 1C**). In contrast, there was no significant difference in the cumulative distribution or average inter-event interval (IEI) between WT and *Shank3* KO groups (**Figure 1D, E**), suggesting that the probability of spontaneous presynaptic glutamate release from the presynaptic site is not affected by the loss of *Shank3*. These findings from miniature recordings suggest that *Shank3* plays a critical role in maintaining postsynaptic glutamatergic receptor function at the MF-CGC synapse.

### Deficiency of *Shank3* alters the biophysical properties of AMPAR and does not affect the presynaptic function

Next, we investigated the change in MF-CGC synapse in *Shank3* KO mice by electrically stimulating the presynaptic terminal and recording the evoked EPSCs (eEPSCs) from CGCs in a whole-cell voltage clamp configuration at -60 mV in the presence of gabazine (**Figure 2A**). As we increased the stimulus intensity from 50 to 300 µA, we observed a modest increase in the eEPSC amplitude up to 200 µA stimulation in both WT and *Shank3* KO, but with with high variability in the amplitude between individual GCGs (**Figure 2B, C**). The mean amplitude of eEPSC did not significantly differ between genotypes at any given stimulus strength (**Figure 2C**). However, the overall distribution of individual eEPSC events generated from all CGCs indicated a shift toward higher amplitude eEPSCs in the *Shank3* KO compared to the WT group (**Figure 2D, E**), similar to what we observed for CGC sEPSCs previously (Kshetri et al., 2024).

**Figure 2:**
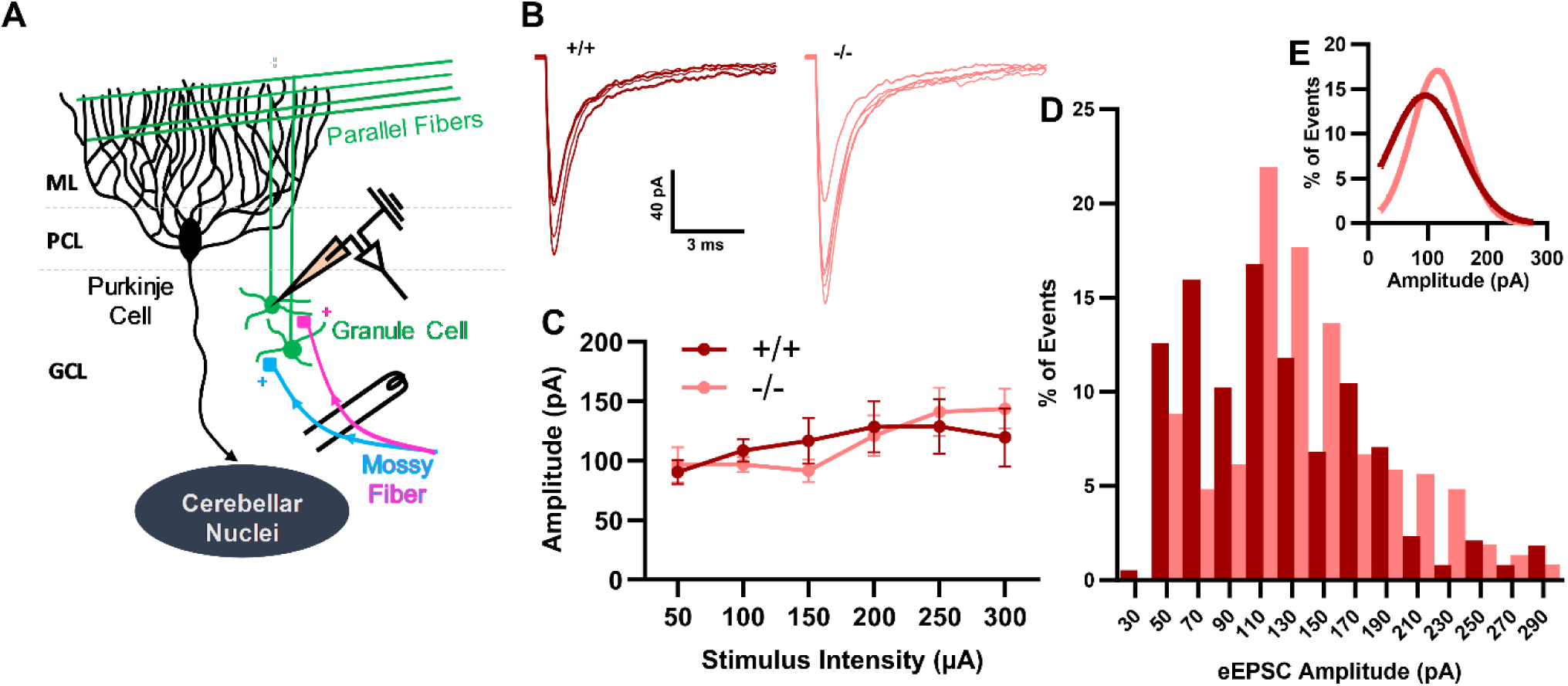
Increased evoked responses at higher stimulation intensities in *Shank3* KO mice. (**A**) Schematic diagram of cerebellar circuit showing the stimulation (mossy fiber, MF) and recording sites (cerebellar granule cell) within the cerebellar cortex. (**B**) Representative traces of eEPSCs at various stimulus intensities in WT (+/+, red) and *Shank3* KO (-/-, light red) mice. (**C**) Quantification of the input-output relationship showing mean eEPSC amplitude ± SEM (bars) at each current intensity applied to stimulate presynaptic MF terminals in WT and *Shank3* KO mice. (**D**) Percentage distribution of amplitude of total eEPSC individual events (excluding failure events) in 20 pA histogram bins from each CGC with the Gaussian fit of the amplitude distribution shown in the inset (**E**). In **C**, WT: n = 40 cells from N = 14 mice; *Shank3* KO: n = 42 cells from N = 20 mice. An unpaired t-test used for parametric and a Mann-Whitney test was used for non-parametric dataset for comparing responses between genotypes at each stimulus intensity. In **D**, WT: n = 11 cells from N = 5 mice, 382 events; *Shank3* KO: n = 13 cells from N = 10 mice, 374 events. Abbreviations: ML: Molecular layer, PCL: Purkinje cell layer, GCL: Granule cell layer.

After stimulating the CGCs at -60 mV to elicit the AMPA response, the holding current was adjusted to +40 mV to record the composite EPSC, with AMPARs generating the fast component and NMDARs producing the slower component. Then, the NMDAR-mediated currents were dissected electrophysiologically by measuring the current value at 15 ms after the stimulus intensity at +40 mV (**Figure 3A**). The average amplitudes of both AMPA- and NMDA-responses at 100 µA stimulation were similar between WT and *Shank3* KO (**Figure 3B, C**). Next, the calculated AMPA/NMDA ratio for CGCs was also similar between the WT and *Shank3* KO groups (**Figure 3D**), which suggests that the basal short-term plasticity at the MF-CGC synapse may not be affected by the loss of *Shank3*.

**Figure 3:**
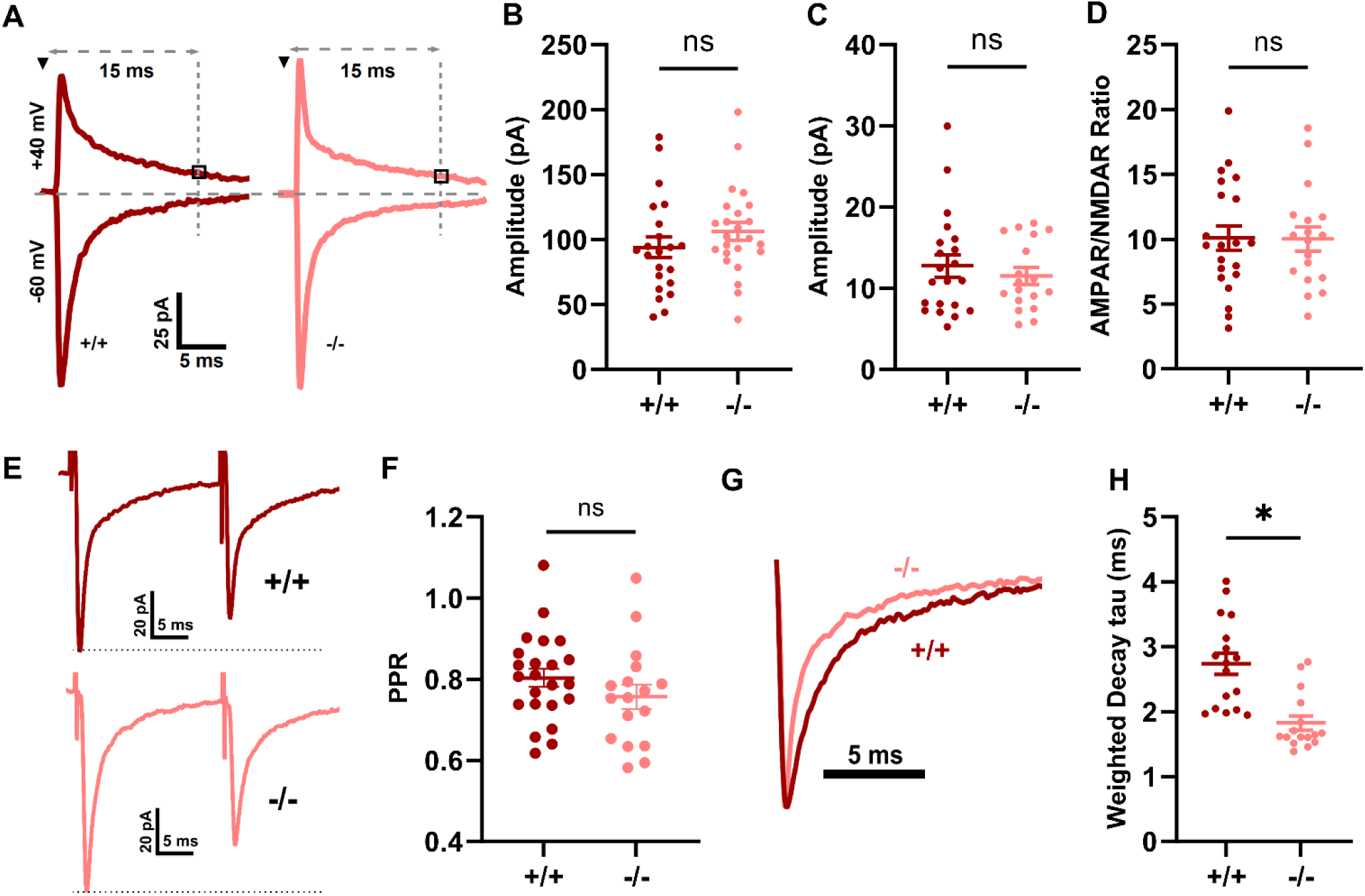
Faster decay kinetics of AMPAR-mediated eEPSC in *Shank3* KO mice. (**A**) Representative traces of AMPAR-mediated eEPSCs at -60 mV holding potential and NMDAR-mediated eEPSCs at +40 mV holding potential in WT and *Shank3* KO mice. (**B**) Average peak amplitudes of AMPAR-mediated eEPSCs. (**C**) Average amplitudes of NMDAR-mediated eEPSCs at +40 mV, 15 ms after stimulation. (**D**) Ratio of average AMPAR to NMDAR amplitudes. (**E**) Representative traces of paired-pulse responses (inter-stimulus interval [ISI]: 20 ms) in WT and *Shank3* KO mice. (**F**) Quantification of the paired-pulse ratio in both genotypes. (**G**) Representative traces of normalized AMPAR-mediated eEPSC illustrating decay kinetics in both genotypes. (**H**) Weighted decay tau values of AMPAR responses evoked by 100 µA stimulation in both genotypes. For panels **B-D, F,** and **H**, individual data points are shown as circles, and bars represent the mean ± SEM. WT: n = 17-23 cells from N = 10-11 mice; *Shank3* KO: n = 16-24 cells from N = 13-18 mice. Statistical significance was determined using an unpaired t-test for **B, D,** and **F**, and a Mann-Whitney test for **C** and **H**. *p < 0.05 indicates a significant difference between WT and KO; ns: not significant.

To determine the role of *Shank3* in the presynaptic function at the MF-CGC synapse, we recorded the paired-pulse ratio (PPR). Our findings showed that the PPR was similar in WT and *Shank3* KO groups (**Figure 3E, F**). The absence of change in PPR and frequency of mEPSC (**Figure 1D, E**) suggest that the absence of *Shank3* does not alter the presynaptic function of the MF-CGC synapse. Interestingly, the evoked AMPAR-mediated response showed significantly faster decay in *Shank3* KO mice than in WT mice (**Figure 3G, H**), indicating the possible change in biophysical properties of AMPARs due to the change in subunit composition or the change in the number of AMPARs at the postsynaptic site.

### Increase in AMPAR-mediated response to uncaged glutamate in *Shank3* KO mice

To determine the AMPAR function irrespective of their presence at the synapse, we supplemented Rubi-Glutamate in the bath and made glutamate available through photolytic cleavage, then recorded the inward currents from CGCs (**Figure 4A**). Glutamate uncaging allows for precise spatial and temporal control of glutamate release, potentially revealing ionotropic glutamate receptor properties that may not be apparent in miniature or evoked EPSC recordings. Brief exposure to blue LED light in the presence of 10 µM gabazine and 0.5 µM TTX while holding a CGC at -70 mV showed a combined AMPA and NMDA response (**Figure 4B**). Furthermore, the NMDA and AMPA components were dissected from the combined response using a pharmacological blocker (10 µM NBQX) and digital subtraction (**Figure 4B**). We observed a significant increase in the combined AMPA- and NMDA-mediated response in *Shank3* KO mice compared to WT (**Figure 4C**). Moreover, upon analyzing the individual AMPA and NMDA currents, we found a significant increase in AMPA current amplitude in the *Shank3* KO group (**Figure 4D**). In contrast, NMDA current amplitudes were similar in both the WT and *Shank3* KO groups (**Figure 4E**). Consequently, the *Shank3* KO group showed an increased AMPA/NMDA ratio compared to the WT (**Figure 4F**). Next, the calculation of current densities for all three currents (AMPA + NMDA, AMPA, and NMDA) demonstrated a significant increase exclusively in AMPAR current density in *Shank3* KO mice (**Figure 4H**). Overall, these findings suggest that the upregulation of AMPAR function at the MF-CGC synapse in *Shank3* KO mice may arise from either an increase in the number of AMPARs at the postsynaptic site or a change in AMPAR subunit composition.

**Figure 4:**
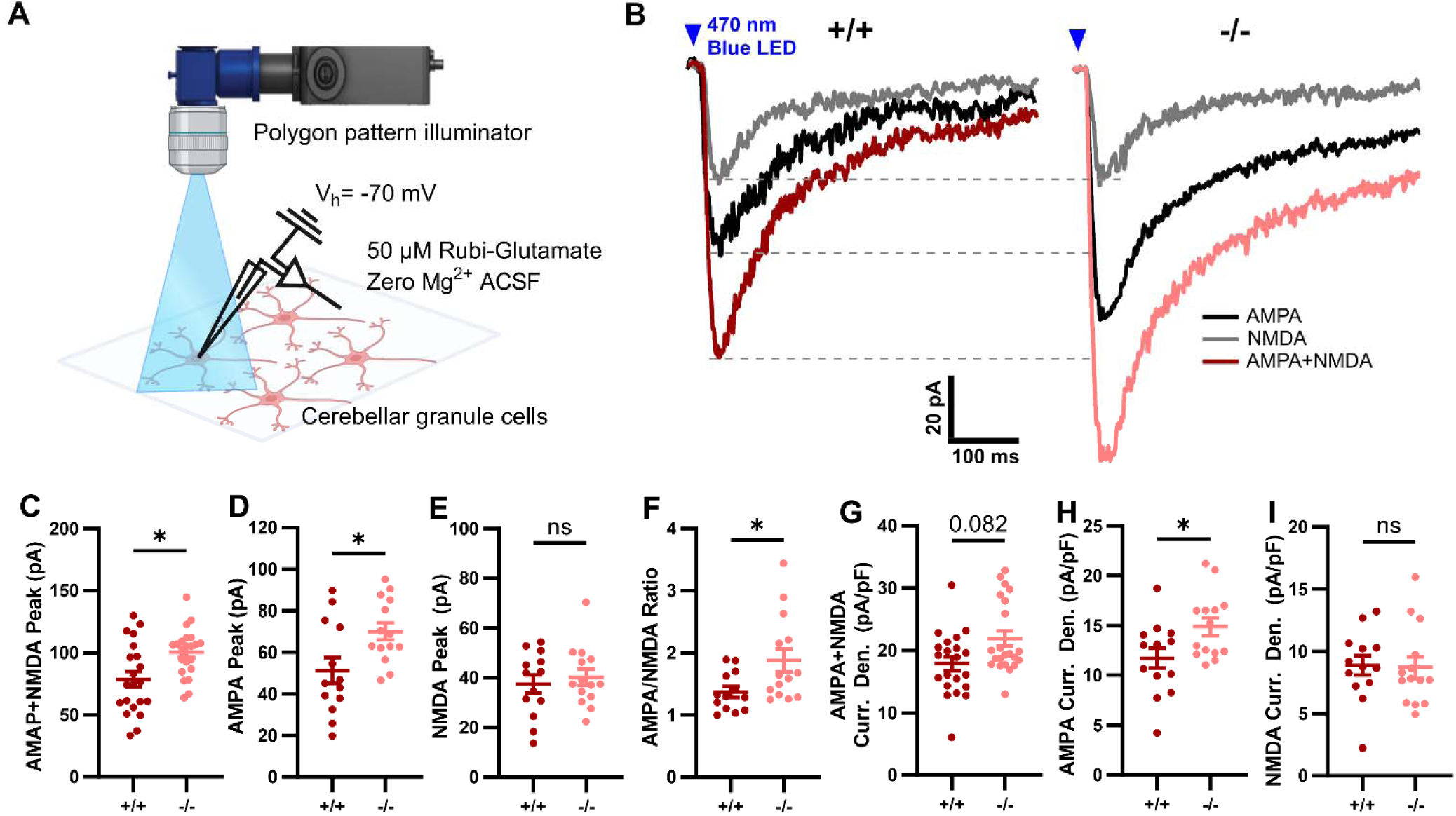
Loss of *Shank3* increases the total AMPAR-mediated response in CGC following glutamate uncaging. (**A**) Schematic diagram of glutamate uncaging experiment, illustrating brief exposure of cerebellar slice to blue LED light for photolytic cleavage of Rubi-glutamate supplied in ACSF (0 mM Mg^2+^) while recording from a CGC at -70 mV. (**B**) Representative current traces of combined AMPA + NMDA (red in WT, light red in *Shank3* KO) current recorded upon light exposure in the presence of Rubi-glutamate, gabazine, and TTX. Subsequent addition of NBQX isolated NMDA current (gray). AMPA current trace (black) was obtained by subtracting NMDA component from the composite AMPA + NMDA current. (**C**) Average peak amplitudes of the combined AMPA + NMDA response. (**D**) Average peak amplitudes of the AMPAR response. (**E**) Average peak amplitudes of the NMDA response. (**F**) Ratio of average AMPAR to NMDAR amplitudes. (**G**) Average current density of combined AMPA + NMDA response. (**H**) Average current density of the AMPA response. (**I**) Average current density of the NMDA response. For panels **C-I**, individual data points are shown as circles, and bars represent the mean ± SEM. WT: n = 13-21 cells from N = 7-8 mice; *Shank3* KO: n = 14-22 cells from N = 6-8 mice. Statistical significance was determined using an unpaired t-test for the data in panels **C-E, H, I** and a Mann-Whitney test for the data in panels **F** and **G**. *p < 0.05 indicates a significant difference between WT and *Shank3* KO; ns: not significant.

### Loss of *Shank3* shows inward rectification and the presence of an increased level of CP-AMPAR in CGCs

AMPAR kinetics are largely determined by the presence or absence of the GluA2 subunit as GluA2-lacking calcium-permeable (CP) AMPARs typically exhibit faster decay kinetics compared to GluA2-containing calcium-impermeable (CI) AMPARs (Anggono & Huganir, 2012; Chojnacka et al., 2023; Cull-Candy & Farrant, 2021; Dolgacheva et al., 2020; C. Guo & Ma, 2021; Henley & Wilkinson, 2016; Hollmann et al., 1991; Traynelis et al., 2010). To confirm the possibility of the presence of CP-AMPARs in CGCs in *Shank3* KO mice, we supplemented spermine in the patch pipette and recorded the eEPSC response at the same setup used for eEPSC recording (**Figure 5A**). The current-voltage curve (I-V curve) showed significant inward rectification at higher potentials (+20, +40, and +60 mV), and a decrease in rectification index (RI) value in *Shank3* KO cells than the WT cells, which indicates a relative incrase in the presence of CP-AMPARs in CGCs in *Shank3* KO mice (**Figure 5B, C, D**).

**Figure 5:**
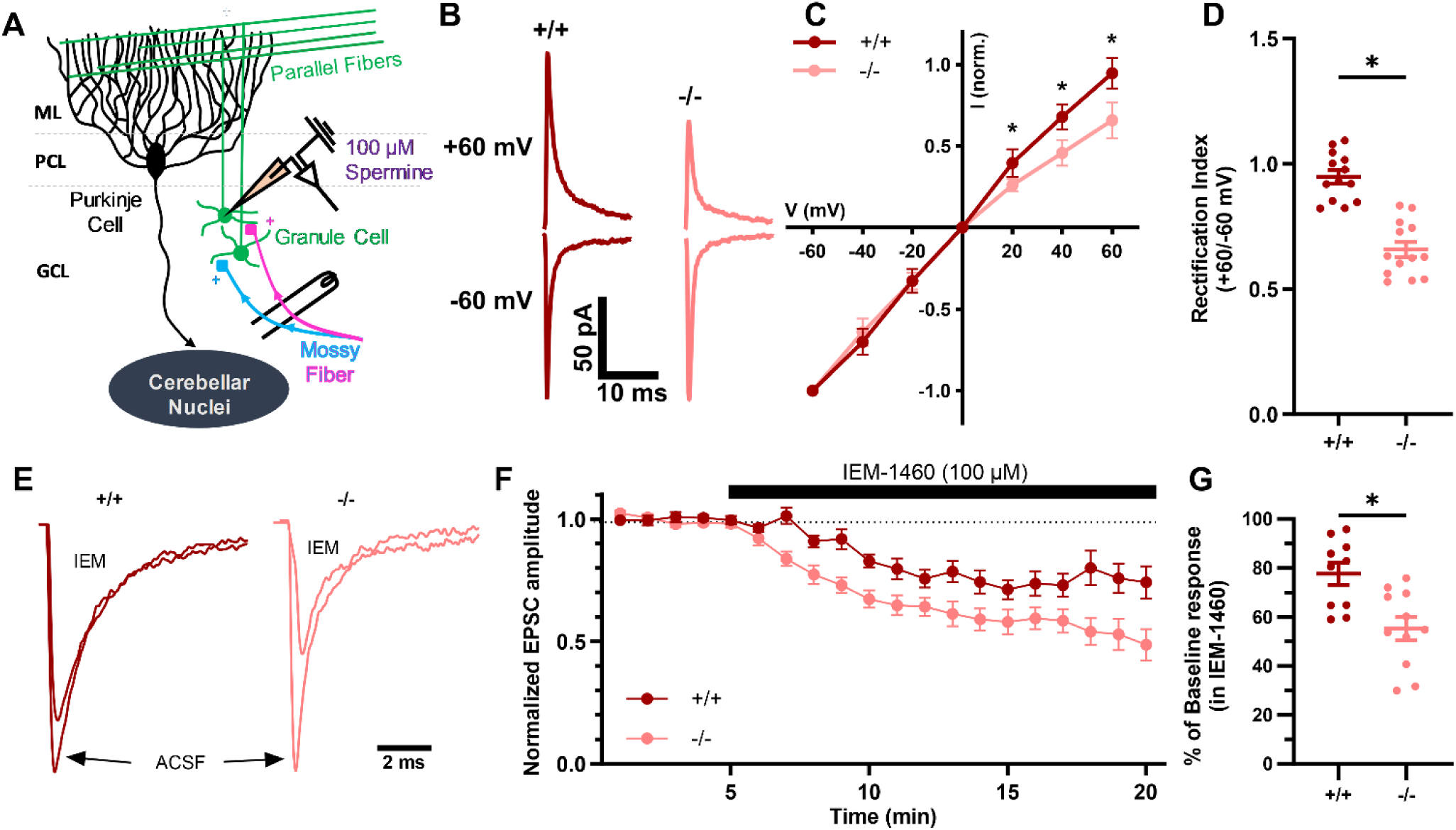
CGCs show inward rectification and an increased proportion of CP-AMPARs in *Shank3* KO mice. (**A**) Schematic diagram of the cerebellar circuit, illustrating the stimulation of MFs and recording a CGC in the presence of intracellular spermine. (**B**) Representative current traces of AMPAR-mediated response from CGCs of WT and *Shank3* KO at -60 and +60 mV. (**C**) Normalized current-voltage (I-V) graph showing the eEPSC amplitude. Data points represent the mean ± SEM, normalized to the current at -60 mV. (**D**) Rectification index values for CGCs from WT and *Shank3* KO. (**E, F**) Example (**E**) and group average (**F**) AMPAR-mediated EPSC responses before (ACSF) and during the IEM-1460 application, normalized to the average baseline response recorded over 5 min before drug application. (**G**) Percentage of baseline response calculated from the average of the last 3 min of recording in IEM-1460 from both genotypes. In panels **D** and **G**, individual data points are shown as circles, and bars represent the mean ± SEM. WT: n = 10-13 cells from N = 6-7 mice; *Shank3* KO: n = 11-13 cells from N = 3-7 mice. Statistical significance was determined using an unpaired t-test for the data in panels **B, D,** and **F**. *p < 0.05 indicates a significant difference between WT and *Shank3* KO. Abbreviations: ML: Molecular layer, PCL: Purkinje cell layer, GCL: Granule cell layer

Next, we further determined the presence of increased CP-AMPARs by applying IEM-1460, a CP-AMPAR blocker (Cull-Candy & Farrant, 2021; da Silva & Schröder, 2023; Suyama et al., 2017; Twomey et al., 2018). After the application of IEM-1460, AMPAR-mediated currents decreased relative to the baseline in WT neurons, suggesting the presence of CP-AMPARs in CGCs (**Figure 5E-G**). These findings align with previous studies that have shown the presence of CP-AMPARs in CGCs (Hack & Balázs, 1995; Savidge & Bristow, 1997). When comparing the degree of IEM-1460-induced reduction in the AMPA response between WT and *Shank3* KO neurons, as predicted, we observed a significantly greater decrease in the AMPA response in *Shank3* KO neurons compared to WT (**Figure 5E-G**). The enhanced sensitivity to IEM-1460 in *Shank3* KO further supports the conclusion that the MF-CGC synapse has an elevated proportion of GluA2-lacking CP-AMPARs.

Since our electrophysiology data suggest an increase in the level of CP-AMPARs in the CGCs of the *Shank3* KO mice, which may lead to an increase in excitability, resulting in excitotoxicity and neuronal loss. Therefore, to determine the possibility of neuronal loss due to the loss of *Shank3*, we stained cerebellar slices with a neuronal marker, NeuN and counterstained with a nuclear marker, Hoechst (**Supplementary Fig. 2A-F**). Since most of the neurons present in the cerebellar cortex are CGCs, counting the NeuN-positive cells will determine if neuronal loss occurred due to the loss of *Shank3*. Following the quantification, we did not find any difference in the number of NeuN-positive cells when comparing the WT and *Shank3* KO groups at the adult time point (4-6 months) (**Supplementary Fig. 2G**). This result suggests that the loss of *Shank3* mainly affects synaptic transmission at the MF-CGC synapse but does not lead to excitotoxicity.

### Reduced IBA1-stained microglial surface area in *Shank3* KO mice

We investigated changes in microglial morphology and number in *Shank3* KO mice using immunostaining with ionized calcium binding adaptor molecule 1 (IBA1) (**Figure 6A, B**). Following staining, we quantified the surface area covered by IBA1-stained microglia (**Figure 6A′, B′**). The total surface area covered by IBA1-stained microglia was significantly reduced in the *Shank3* KO group relative to the WT controls (**Figure 6C**). Furthermore, the distribution plot of the surface area measurement showed that the majority of microglia in the *Shank3* KO group covered a smaller surface area relative to the WT group (**Figure 6E**), indicating a shift toward a less ramified morphology. However, the number of microglia in the cerebellar cortex was similar between WT and *Shank3* KO mice (**Figure 6D**).

**Figure 6:**
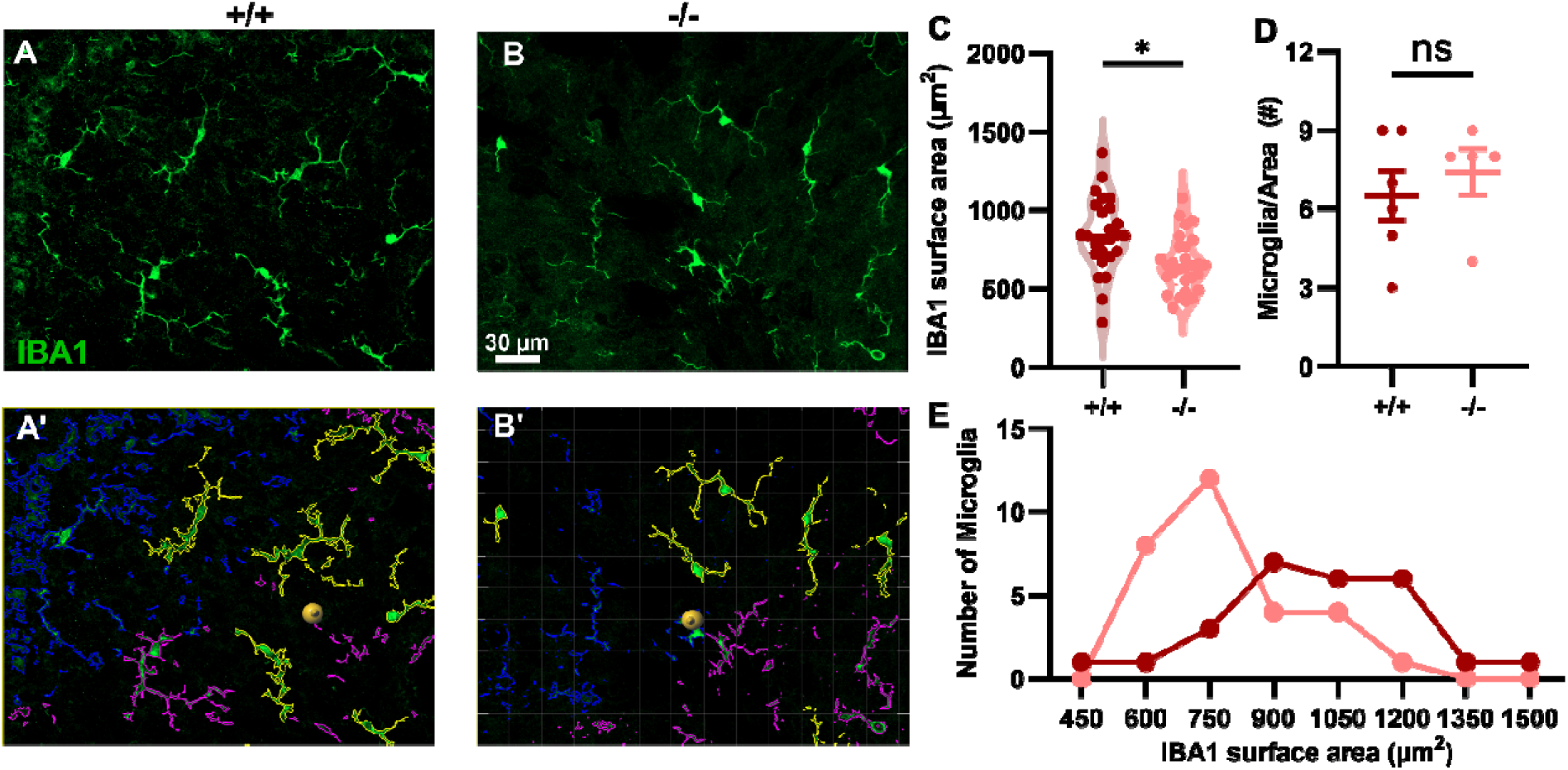
*Shank3* KO mice showed a reduced IBA1-stained fluorescence area in the cerebellum. (**A, B**) Representative images of immunolabeled microglia with IBA1 cerebellar granule cell layer of sagittal cerebellar sections from WT (+/+) and *Shank3* KO (-/-) mice. (**A**_′_**, B**_′_) Corresponding images showing the surface area occupied by IBA1-positive microglia (outlined in yellow and magenta) from the cerebellar granule cell layer presented in panels **A** and **B**, respectively. (**C**) Violin plot of the quantification of surface area covered by individual IBA1-stained microglia in WT and *Shank3* KO mice. Individual data points represent the surface area of each IBA1-positive microglial cell; bars represent the median. (**D**) Quantification of the total number of microglia per image field (265 × 265 µm) in WT and *Shank3* KO mice. Individual data points represent values from each image field, and bars indicate mean ± SEM. (**E**) A histogram showing the frequency distribution of the total surface area covered by IBA1-stained microglia in WT and *Shank3* KO mice. A total of 5-6 images were analyzed per genotype (26-29 microglia per genotype). N = 3-5 mice/genotype. Statistical significance was assessed using unpaired t-tests in panels **C** and **E**. * indicates p < 0.05; ns: not significant.

To determine the presence of astrocyte reactivity in germline *Shank3* KO mice, we also performed immunostaining using the astrocytic marker Glial Fibrillary Acidic Protein (GFAP) along with SRY-box transcription factor 9 (SOX9), a nuclear marker for astrocytes (**Supplementary Fig. 1A-D**). The percentage area covered by GFAP fluorescence was comparable between WT and *Shank3* KO mice (**Supplementary Fig. 1E**). Similarly, no significant difference was observed in GFAP fluorescence intensity between the two groups (**Supplementary Fig. 1F**). Additionally, staining with SOX9 revealed no difference in the number of SOX9-positive astrocytes between the WT and *Shank3* KO groups (**Supplementary Fig. 1G**). Together, these results indicate that the absence of *Shank3* does not induce astrocyte reactivity or alter astrocyte number in the cerebellar cortex; however, the presence of less ramified microglia suggests a possible shift toward an activated microglial state.

## Discussion

In this study, we investigated the role of *Shank3* in regulating synaptic function at the MF-CGC synapse, a critical part of the cerebellar circuit that integrates sensory-motor information in the cerebellar cortex. Additionally, we also studied the alteration in astrocyte reactivity and microglia morphology in *Shank3* KO mice. Our electrophysiological and immunofluorescence data demonstrate the role of *Shank3* in maintaining AMPAR function and subunit composition at the postsynaptic site while indicating that the presynaptic release property remains unaffected by its absence. Additionally, less ramified microglia were observed in *Shank3* KO mice. This change in synaptic function, altering the excitatory neurotransmission in CGCs, and altered microglia sheds light on the potential mechanism of cerebellum involvement in the development of ASD-like phenotypes in *Shank3* KO mice. One of the main findings of our electrophysiology experiments is the enhancement of postsynaptic AMPAR-mediated response in CGC from *Shank3* KO mice compared to WT controls. First, we observed the enhancement of amplitude but no change in the frequency of mEPSC in adult *Shank3*^Δ*ex4-22*^ mice (**Figure 1A-E**). However, studies performed across various brain regions and using different versions of *Shank3* mutant mice have shown inconsistent results regarding mEPSC parameters. For instance, *Shank3B* KO mice showed a reduction in both frequency and amplitude of mEPSC in dorsolateral striatal medium spiny neurons (MSNs) (Peça et al., 2011) and anterior cingulate cortex (ACC) pyramidal neurons (B. Guo et al., 2019). However, Guo et al. reported a decrease in mEPSC amplitude without a change in frequency in thalamocortical cells of *Shank3B* KO mice (B. Guo et al., 2024). Furthermore, studies using *Shank3*^Δ*14–16*^ mouse model reported an increase in frequency but no change in amplitude in mPFC layer 2/3 neurons (Yoo et al., 2019), whereas both frequency and amplitude were reduced in the dorsolateral striatum (Yoo et al., 2018). Additionally, the mouse model of frameshift mutation (InsG3680 mutation) in *Shank3* showed a significant increase in mEPSC amplitude but no change in frequency in the dorsal striatum (Zhou, Kaiser, Monteiro, Zhang, Van der Goes, et al., 2016). Similarly, *Shank3* knockdown in the anteromedial bed nucleus of the stria terminalis (BNST) neurons resulted in increased mEPSC amplitude and decreased frequency (Contestabile et al., 2023). These discrepancies suggest the brain-region-specific and/or mutation-specific alterations of the *Shank3* isoform across the brain regions. Among these, CGCs may show a unique response to the loss of *Shank3* compared to other brain regions.

Additionally, we found an increase in AMPAR response and AMPA/NMDA ratio in the uncaging experiment, which further supports the idea of enhancement of AMPAR function (**Figure 4D, F**), either due to an increase in expression or the biophysical properties of AMPARs. Together, these increases in AMPAR-response observed in CGCs in our study could arise from several potential factors. Shank3 is known to be involved in the trafficking and synaptic localization of AMPARs (H. T. T. Ha et al., 2018; Yang et al., 2024). Therefore, loss of *Shank3* may lead to the alteration of AMPAR subunit composition, potentially incorporating subunits that show higher conductance. Another possibility of an increase in mEPSC amplitude could be an increased insertion of functional AMPARs at the postsynaptic membrane (Chen et al., 1999; Glasgow et al., 2019), however, some evidence from other brain regions suggests *Shank3* deficiency may reduce AMPAR function (Xu et al., 2024; Xue et al., 2024). Beyond AMPAR subunit composition, their function is also modulated by integral auxiliary subunits, including Transmembrane AMPAR Regulatory Proteins (TARPs), Cornichon homologs (CNIHs), Cysteine-knot AMPAR Modulating Proteins (CKAMPs), and Germ cell-specific gene 1-like protein (GSG1L) (Anggono & Huganir, 2012; Bissen et al., 2019; Cull-Candy & Farrant, 2021; Qneibi et al., 2024). These auxiliary subunits shape AMPAR biophysical properties by dynamically tuning channel gating kinetics, single-channel conductance, and sensitivity to polyamine block, thereby fundamentally determining the characteristics of synaptic transmission and plasticity (Coombs et al., 2023; Cull-Candy & Farrant, 2021; Jacobi et al., 2021; Kamalova et al., 2020; Miguez-Cabello et al., 2025; Qneibi et al., 2024; Rozov et al., 2018). Disruption of *Shank3* could indirectly affect the interaction between AMPARs and their auxiliary subunits, potentially enhancing AMPAR function. While stable mEPSC amplitudes were reported during LTP at the mossy fiber-granule cell synapse (Sola et al., 2004), a study using Islet Brain-2 (IB2, a gene commonly deleted in PMS) KO found an increase in quantum size in CGCs during LTP (Soda et al., 2019). Therefore, if the LTP mechanism is altered at this synapse in *Shank3* KO mice, it could account for the increased postsynaptic receptor function observed in our study.

The absence of change in mEPSC frequency in our study suggests that the presynaptic neurotransmitter release machinery remains intact at the MF-CGC synapse (**Figure 1D, E**). This finding is consistent with some studies performed in different brain regions with *Shank3* mutations mentioned above in the discussion (B. Guo et al., 2024; Jaramillo et al., 2016; X. Wang et al., 2011; Zhou, Kaiser, Monteiro, Zhang, Van der Goes, et al., 2016) (Zhou, Kaiser, Monteiro, Zhang, Van der Goes, et al., 2016), suggesting that *Shank3* deficiency does not universally impact the probability of spontaneous presynaptic glutamate release. Conversely, a number of studies have reported either an increase (Yoo et al., 2019) or decrease (B. Guo et al., 2019; Kouser et al., 2013; Peça et al., 2011; W. Wang et al., 2017; Yoo et al., 2018) in mEPSC frequency, indicating that the presynaptic effects of *Shank3* loss may depend on the brain region, the specific *Shank3* isoform affected, and the developmental stage.

We observed a shift toward higher amplitude AMPAR-mediated eEPSCs (**Figure 2D, E**) without a corresponding change in AMPA/NMDA ratio in the *Shank3* KO group (**Figure 3D**). The stability in the AMPA/NMDA ratio aligns with some previous findings (Peça et al., 2011; Zhou, Kaiser, Monteiro, Zhang, Van der Goes, et al., 2016) but contrasts with the others. For instance, a study showed a reduction in both the eEPSC input-output curve relationship and the AMPA/NMDA ratio in ACC pyramidal neurons of *Shank3* InsG3680 mutant mice (B. Guo et al., 2019). Variability regarding changes in the AMPA/NMDA ratio has been documented in the literature for *Shank3* KO models. Consistent with our results, some studies have observed no change in the AMPA/NMDA ratio (Peça et al., 2011; Zhou, Kaiser, Monteiro, Zhang, Van der Goes, et al., 2016), whereas others have shown a decrease in the NMDA/AMPA ratio (Jaramillo et al., 2016; Kouser et al., 2013; Speed et al., 2015). Interestingly, Jaramillo et al. (2016) showed no alterations in the NMDA/AMPA ratio in the hippocampus of *Shank3*^Δ*e4-9*^ mutant mice; however, the same study observed a significant reduction in the NMDA/AMPA ratio in the striatum (Jaramillo et al., 2016), which again highlights that the effects of *Shank3* mutations on synaptic function can be brain region-specific. In this context, our findings suggest that the cerebellum, with its distinct developmental timeline and circuit organization, may exhibit unique synaptic responses to *Shank3* deficiency.

Multiple lines of evidence from our study indicate that the loss of *Shank3* may not alter the probability of neurotransmitter release at the MF-CGC synapse. We found that both mEPSC frequency (**Figure 1D, E**) and PPR (**Figure 3F**) were similar between WT and *Shank3* KO, suggesting that the synaptic deficits observed in our model may be predominantly postsynaptic in origin. These findings are consistent with several previous studies that reported no significant changes in PPR in various *Shank3*-deficient mouse models (B. Guo et al., 2019; Jaramillo et al., 2016, 2017; Kouser et al., 2013; Moutin et al., 2021; Peça et al., 2011; Speed et al., 2015; X. Wang et al., 2011; Zhou, Kaiser, Monteiro, Zhang, Van der Goes, et al., 2016). However, Wang et al. (2017) noted a significant increase in PPR exclusively in the D2 MSNs (striato-pallidal indirect pathway) of the *Shank3B* KO mice, indicating a reduced presynaptic release probability specifically at the glutamatergic terminals innervating D2 MSNs, whereas PPR in D1 MSNs (striato-nigral direct pathway) was similar between WT and *Shank3* KO (W. Wang et al., 2017). Although most studies collectively suggest that *Shank3* loss does not generally alter the presynaptic release probability, it is possible that *Shank3* might form distinct signaling complexes or express specific isoforms with unique functions in different cell types or brain regions (W. Wang et al., 2017).

Our observation of a fast decay in AMPAR-mediated evoked EPSCs in CGCs (**Figure 3G, H**) of *Shank3* KO mice suggests a possible change in subunit composition in AMPARs, offering a novel perspective on the role of *Shank3* in regulating glutamatergic transmission within the cerebellum. The functional diversity of AMPARs is mainly determined by their subunit composition (Anggono & Huganir, 2012; Chojnacka et al., 2023; Cull-Candy & Farrant, 2021; Dolgacheva et al., 2020; C. Guo & Ma, 2021; Henley & Wilkinson, 2016; Hollmann et al., 1991; Traynelis et al., 2010). The absence of the GluA2 subunit has a profound effect by causing calcium permeability, higher single-channel conductance, faster kinetics, and inward rectification in its current-voltage relationship due to voltage-dependent block by intracellular polyamines (Burnashev et al., 1992; Cull-Candy & Farrant, 2021). In addition to faster decay kinetics, we have noticed inward rectification and a comparatively high CP-AMPAR response in *Shank3* KO, suggesting *Shank3* loss may affect the homeostatic balance of surface expression of CP- and CI-AMPARs in the CGCs. The change in proportion of inwardly rectifying CP-AMPAR could confound the measurement of amplitude at higher potentials (Kauer & Malenka, 2007), which might explain the similar AMPA/NMDA ratio in eEPSC response in both WT and *Shank3* KO group.

In the majority of neurons across various brain regions, the CP-AMPARs are rapidly replaced with CI-AMPARs after P14 in rodents (Diering & Huganir, 2018). Concurrently, the expression of all *Shank3* isoforms significantly increases after P7, reaching a peak around four weeks postnatally (X. Wang et al., 2014). A significant mechanism underlying synaptic plasticity involves activity-dependent switch from CP-AMPARs to CI-AMPARs (S. J. Liu & Savtchouk, 2012; S. J. Liu & Zukin, 2007). Synaptic CP-AMPARs show a unique self-regulating feature, whereby repetitive activation of these receptors triggers their replacement by GluA2-containing CI-AMPARs. This phenomenon was initially observed and well-characterized at the parallel fiber to stellate cell synapse in the cerebellum (S.-Q. J. Liu & Cull-Candy, 2000).

Shank3, along with other postsynaptic proteins and zinc, has been shown to interact and assist in the trafficking of GluA2-containing AMPARs at the synaptic membrane (Bariselli et al., 2016; H. T. T. Ha et al., 2018; Sheng & Kim, 2000). This is mediated through the interaction of Shank3 with Glutamate Receptor Interacting Proteins (GRIPs) through its SH3 domain (Bozdagi et al., 2010; Manning et al., 2024; Sala et al., 2015; Sheng & Kim, 2000). Furthermore, PDZ domain-containing proteins like GRIPs (Dong et al., 1997, 1999; S. J. Liu & Cull-Candy, 2005) and protein interacting with C kinase (PICK1) (Xia et al., 1999), interact with the C-terminal of the GluA2 subunit. This molecular bridge, formed through the interactions among Shank3, PDZ-domain containing proteins (GRIPs and PICK1), and GluA2, is posited to be a crucial mechanism for the appropriate synaptic localization and stabilization of GluA2-containing CI-AMPARs. The molecular mechanisms underlying the switch from CP- to CI-AMPARs involve an increase in intracellular Ca^2+^ through CP-AMPARs. Elevated intracellular Ca^2+^ subsequently activates protein kinase C (PKC), which in turn activates PICK1 (S. J. Liu & Savtchouk, 2012; S. J. Liu & Zukin, 2007). Concurrently, PKC may phosphorylate GluA3, disrupting the interaction between GluA2-lacking AMPARs and GRIP, leading to the loss of these CP-AMPARs from the synapse (Gardner et al., 2005; S. J. Liu & Cull-Candy, 2005). Therefore, in the case of *Shank3* KO, the loss of Shank3 may interrupt the AMPAR trafficking, affecting both the insertion of CI-AMPAR into the membrane and the internalization of CP-AMPAR. Thus, the synaptic level of CP-AMPAR may remain high, resembling that of the early postnatal stage, potentially impairing the maturation of MF-CGC synapses. Additionally, the elevated intracellular Ca^2+^ level could trigger intracellular signaling cascades, altering intrinsic neuronal excitability and influencing cerebellar circuit output (S. J. Liu & Savtchouk, 2012; S. J. Liu & Zukin, 2007). Therefore, the dysregulation of CP- and CI-AMPARs dynamics at MF-CGC synapse in *Shank3* KO mice may impair both synaptic transmission and neuronal excitability output, ultimately contributing to cerebellar-related behavioral deficits.

Our observation of reduced microglial processes, suggestive of an activated state, provides preliminary evidence for potential microglial involvement in the cerebellar pathology of *Shank3* KO mice. In contrast, a study using *Shank3* heterozygous mice reported no differences in IBA1-positive microglia density or cell body size in the hippocampus, mPFC, or striatum (Cope et al., 2016), suggesting that gross IBA1 morphology and cell body size might not be consistently altered in *Shank3* deficiency across all brain regions or developmental time points. Microglial morphology has been used to indicate their functional state. The highly ramified cells are thought to represent a ’resting’ or homeostatic phenotype, and cells exhibiting retracted processes and enlarged soma are considered ‘activated’ (Davis et al., 2017; Morgan et al., 2010; Savage et al., 2019). However, recent perspectives highlight heterogeneity of microglial states and emphasize that morphology analysis itself does not provide insights into their functional capabilities (J. Kim et al., 2023; Paolicelli et al., 2022). Therefore, our findings of IBA1 morphology in *Shank3* KO microglia should be carefully interpreted.

## Conclusion

Several studies have explored the role of *Shank3* in various forebrain regions, but its function in the cerebellum remains underexplored, despite its involvement in both motor and multiple non-motor processes. By using various electrophysiological methods, we have found that the loss of *Shank3* selectively disrupts the postsynaptic receptor function without impacting the presynaptic glutamate release at the MF-CGC synapse. Furthermore, we have found that the loss of *Shank3* results in upregulation of AMPAR-mediated response and a shift towards a high level of CP-AMPARs at the MF-CGC synapse. These findings indicate an impairment in synapse maturation at the MF-CGC synapse, potentially leading to a change in excitability and synaptic plasticity within the cerebellar cortex. Moreover, the less ramified microglial processes in *Shank3* KO suggested the possibility of activated microglia in the cerebellar cortex.

Given that the present study was conducted using the germline *Shank3* KO mouse model, future studies using the most appropriate conditional CGC-specific *Shank3* KO model may address the cell-specific and developmental role of *Shank3* in CGCs. Additionally, targeted pharmacological or genetic interventions to restore CI-AMPAR expression may provide insights into the therapeutic potential for addressing AMPAR dysfunction in ASD.

Together, our findings provide novel insights into the possible role of *Shank3* in regulating the balance between the dynamic regulation of CP- and CI-AMPARs at the MF-CGC synapse. This reveals a possible novel mechanism by which the loss of *Shank3* may disrupt this homeostatic balance, altering the cerebellar circuitry. This study contributes significantly to our understanding of the role of *Shank3* in the cerebellar cortex and the neurobiological underpinnings of *Shank3*-associated cerebellar-related behavioral deficits.

## Declarations

### Ethics approval

All procedures involving animals were performed in accordance with protocols approved by the Institutional Animal Care and Use Committee at Southern Illinois University – School of Medicine.

### Consent for publication

Not applicable.

### Availability of data and materials

The datasets used and/or analyzed during the current study are available from the corresponding author on reasonable request with statistical analysis results included with this published article’s supplementary information files.

### Authors’ contributions

RK drafted the manuscript with BDR. RK collected and analyzed the electrophysiology and immunofluorescence experiments data. BDR performed final review of data analysis and prepared data figures with RK. BDR conceived of the project with RK and BDR oversaw the data acquisition and generation of the final manuscript.

## Acknowledgements

We would like to thank the National Institute of Mental Health for providing funding for this project.

