## Supplementary Data for "*Shank3* establishes AMPA receptor subunit composition at cerebellar mossy fiber-granule cell synapses and shapes regional microglia activation"

**Supplementary 2**

**Methods**

**GFAP and SOX9 staining**

The brain slices were initially placed in 0.01M sodium citrate buffer (pH 6.0), and antigen retrieval was performed at 75 °C for 15 min. The brain slices were then permeabilized in a solution containing 95% methanol and 5% acetic acid for 10 min, followed by IHC/ICC Blocking Buffer (eBiosciences) with 0.5% Triton X-100 for 1 h. Next, the slices were incubated in primary antibodies chicken IgG anti-GFAP (1:1000; Novus Biologicals, catalog# NBP1-05198) and rabbit IgG anti-SOX9 (1:1000; EMD Millipore, catalog# AB5535) overnight at 4 °C. Slices were then washed with PBS and incubated in secondary antibodies goat IgG anti-chicken Alexa Fluor 488 (1:500; Thermo Fisher Scientific, catalog# A11039) and donkey IgG anti-rabbit DyLight 594 (1:500; Novus Biologicals, catalog# NBP1-75637). The nuclei were counterstained with Hoechst 33342 (1:2000, Thermo Fisher Scientific, catalog# H3570). Finally, tissue sections were washed and transferred to glass slides and mounted with Prolong Gold (Invitrogen). Imaging of the inner granule cell layer of cerebellar lobe 4 &5 from the stained brain slices was performed by capturing Z-stack images containing 7 optical slices at an interval of 1 µm, using a 40X oil immersion objective (numerical aperture 1.3) on a confocal microscopy system (Zeiss LSM800) at a resolution of 1,024 × 1,024 pixels.

*Data analysis:* First, the orthogonal projections of Z-stack images were generated using ZEN Lite software (v 3.7 edition; Zeiss). To determine GFAP fluorescence intensity, a single channel image (in JPG format) was exported from ZEN Lite and converted into an 8-bit image in ImageJ/ Fiji (NIH). The image background was subtracted in ImageJ using the default settings (rolling ball radius 50 pixels). GFAP fluorescence intensity was then measured across the entire image. To quantify background fluorescence, a 10 × 10 µm2 square box was placed in the darkest area of the image, and the background intensity was measured within this region. The background intensity was subtracted from the previously measured GFAP fluorescence intensity to obtain the actual fluorescence intensity in the image. The number of astrocytes was determined by counting all SOX9-positive astrocyte cell bodies in the orthogonal projection image. Two images from each animal were analyzed, with 5 animals per genotype.

**NeuN staining for CGC density**

The brain slices were initially placed in 0.01M sodium citrate buffer (pH 6.0), and antigen retrieval was performed at 75 °C for 30 min. The brain slices were then permeabilized in 0.25% Triton X-100 in PBS for 30 min, followed by blocking in IHC/ICC Blocking Buffer (eBiosciences) containing 0.25% Triton X-100 for 1 h. Subsequently, the slices were incubated in primary antibody rabbit IgG anti-NeuN (1:1000, Novus Biologicals, catalog# NBP1-92716) overnight at 4 °C. Slices were then washed with PBS and incubated in secondary antibody donkey IgG anti-rabbit DyLight 488 (1:1000; Thermo Fisher Scientific, catalog# SA5-10038). The nuclei were counterstained with Hoechst 33342 (1:2000, Thermo Fisher Scientific, catalog# H3570). Finally, tissue sections were washed and transferred to glass slides and mounted with Prolong Gold (Invitrogen). Imaging of NeuN-positive CGCs in the inner granule cell layer of cerebellar lobe 4 &5 from the stained brain slices was performed by capturing Z-stack images containing 3 optical slices at an interval of 1 µm using a 40X oil immersion objective (numerical aperture 1.3) on a confocal microscopy system (Zeiss LSM800) at a resolution of 1,024 × 1,024 pixels.

*Data analysis:* The cell density of CGCs was determined by counting the total number of NeuN-positive CGCs within a 50 × 50 µm^2^ square box placed in the inner granule cell layer in cerebellar lobes 4 and 5 using ZEN Lite software (v 3.7 edition; Zeiss). Two images from each animal were analyzed, with five animals per genotype.

**Results**

***Shank3* deletion does not induce astrocyte reactivity**

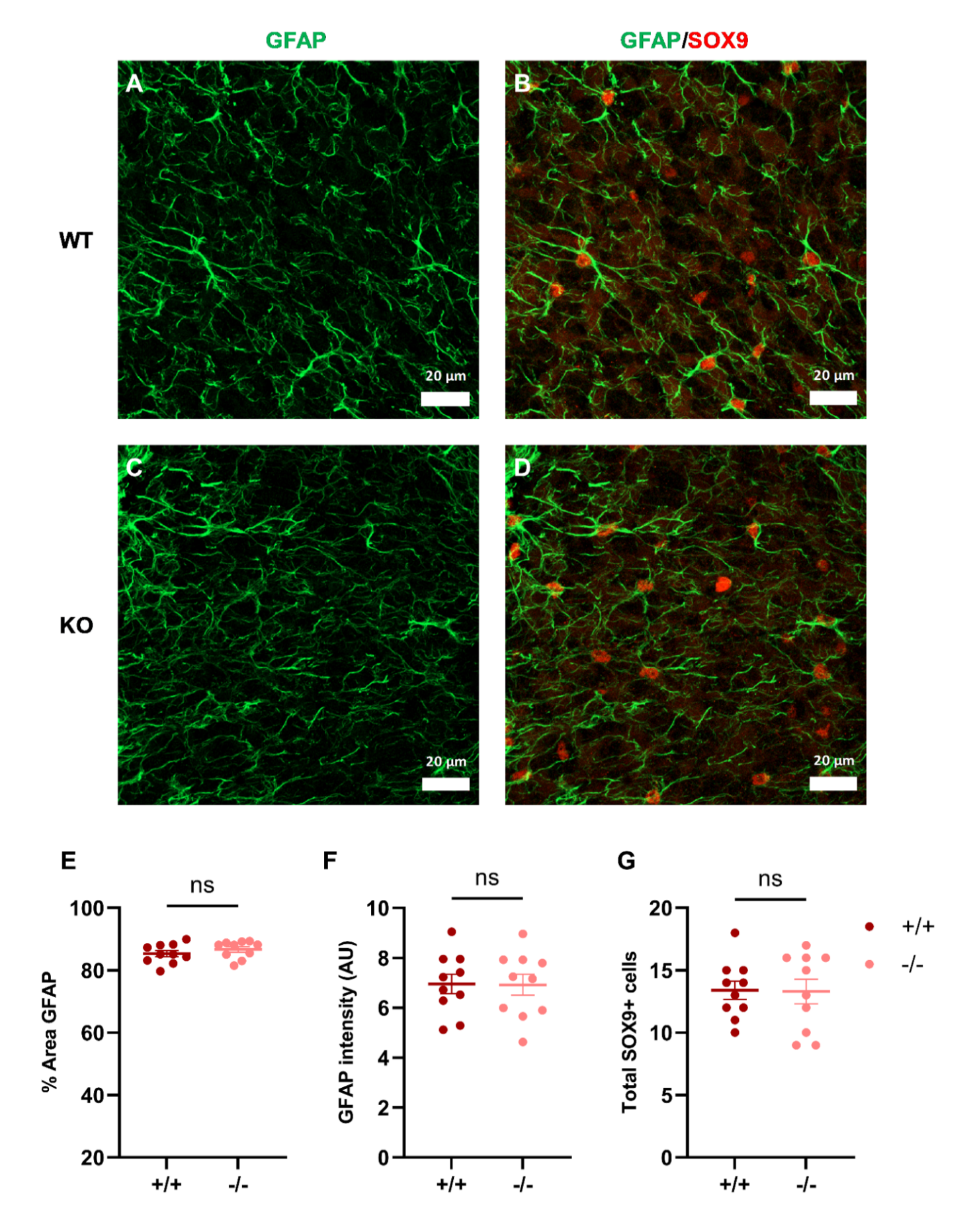

**Supplementary Fig. 1:** **Cerebellar sections immunostained with GFAP and SOX9 in WT and germline *Shank3* KO mice.** (A) A representative WT cerebellum section immunostained for GFAP (green) and (B) both GFAP and SOX9 (red). SOX9 is an astrocyte-specific nuclear marker. (C) A representative *Shank3* KO cerebellum section immunostained for GFAP (green) and (D) both GFAP and SOX9 (red). (E) Quantification of the percentage of area covered by GFAP fluorescence per image field in WT and *Shank3* KO mice. (F) Quantification of GFAP fluorescence intensity in WT and *Shank3* KO mice. (G) Total number of SOX9-positive astrocytes per image field in WT and *Shank3* KO mice. In E-G panels, individual data points represent values from each image field, and bars indicate mean ± SEM. A total of 10 images were analyzed per genotype (N = 5 mice/genotype). Statistical significance was assessed using unpaired t-tests. A p-value < 0.05 was considered statistically significant; ns: not significant.

**Loss of *Shank3* did not cause CGC loss**

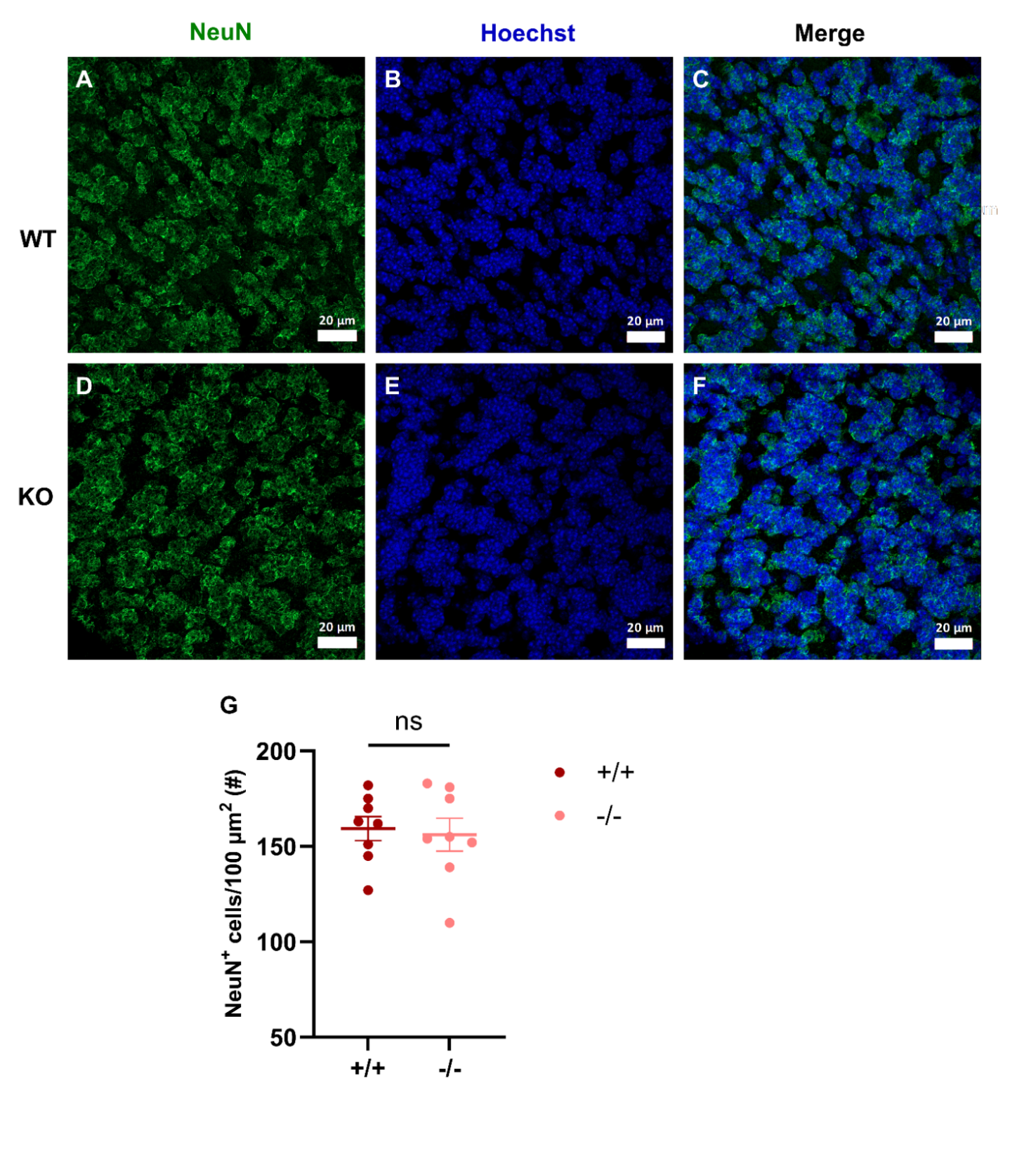

**Supplementary Fig. 2: Cerebellar sections immunostained with neuronal marker (NeuN) in WT and *Shank3* KO mice.** (A-C) Top panel shows a representative image from a WT mouse showing NeuN staining (A), Hoechst (nuclear marker, B), and a merge panel (C). (D-F) Bottom panel shows a representative image from a *Shank3* KO mouse showing NeuN staining (D), Hoechst (nuclear marker, E), and a merge panel (F). (G) Quantification of NeuN-positive cells per 100 µm^2^ area within the cerebellar cortex. Individual data points represent values from each image, and bars indicate mean ± SEM. N = 4 mice/genotype. Two images were analyzed from each mouse. Statistical significance was assessed using unpaired t-tests in panels G. “ns” indicates not significant.
